# Genotype-Dependent Variation in Vitamins B1, B2, B3, B6, B9, and C in Mungbean Sprouts

**DOI:** 10.64898/2026.07.31.742010

**Authors:** Seyoung Jeon, Hakyung Kown, Seung Yeob Song, Sung Don Lim, Jungmin Ha

**Affiliations:** Department of Agriculture, Forestry and Bioresources and Research Institute of Agriculture and Life Sciences, Seoul National University, Seoul 08826, Republic of Korea; Crop Genomics Lab, Plant Genomics and Breeding Institute, Seoul National University, Rm. 4105 Bldg. 200 CALS, 1 Gwanak-ro, Gwanak-gu, Seoul, 08826, Republic of Korea; Food Tech Resources Research Division, Department of Food Sciences, National Institute of Crop and Food Science, Rural Development Administration, Republic of Korea; Molecular Plant Physiology Laboratory, Department of Applied Plant Sciences, Graduate School, Sangji University, Wonju 26339, Republic of Korea

## Abstract

Water-soluble vitamins are essential micronutrients that are not stored extensively in the human body and must therefore be replenished through dietary intake. Mungbean sprouts are widely consumed in some populations; however, their contributions as dietary sources of water-soluble vitamins and the extent of genotype-dependent variations remain insufficiently characterized. In this study, six water-soluble vitamins (B1, B2, B3, B6, B9, and C) were quantitatively and qualitatively analyzed using ultra performance liquid chromatography in sprouts derived from 34 mungbean genotypes. With the exception of pyridoxine (vitamin B6), all analyzed vitamins were detected across all genotypes, and their contents showed significant genotype-dependent variation. To determine whether differences in vitamin composition are reflected in functional responses, antioxidant capacity and enzyme-based bioactivity were evaluated. Antioxidant activity varied significantly among genotypes and showed a positive correlation with nicotinamide (B3) content, while tyrosinase inhibition was positively correlated with thiamine (B1). Evaluation of vitamin contents on a fresh-weight basis demonstrated that a single 104 g serving of mungbean sprouts can provide 46%– 206% of the recommended daily allowance of thiamine depending on genotype, while supplying all analyzed water-soluble vitamins, including B1, B2, B3, B6, B9, and C. This study’s findings support the potential of mungbean sprouts to serve as plant-based dietary sources of water-soluble vitamins and offer a basis for selecting genetic resources to improve the vitamin-related nutritional quality of mungbean sprouts.

## Introduction

Legumes generally contain 20%–45% protein on a dry weight basis and are major dietary sources of protein in developing countries, where access to animal protein is often limited [1], [2], [3]. Legumes are capable of symbiotic nitrogen fixation, allowing their cultivation under low-fertility soil conditions with reduced nitrogen fertilizer inputs [4]. Due to these agronomic and nutritional characteristics, legumes are widely cultivated in developing countries, where they are important components of daily diets [5], [6].

In many developing countries, diets rely heavily on a limited number of staple foods, including cereals and legumes [7]. As a result, the populations of such countries reportedly often experience deficiencies in one or more essential micronutrients, including vitamins [8]. These deficiencies are largely attributed to staple crop–based diets low in vitamin-rich fruits, which are associated with relatively high costs, perishability, and limited economic accessibility [9]. Nutritional-deficiency diseases such as beriberi (associated with low levels of thiamine) and pellagra (nicotinamide), as well as anemia and neurological disorders (folate) and scurvy (ascorbic acid), remain important public health concerns [10].

Vitamins are essential for cellular metabolism and physiological functions. Among them, water-soluble vitamins are not stored extensively in the human body and are readily excreted, necessitating continuous dietary replenishment to maintain metabolic homeostasis [11]. Water-soluble vitamins, including thiamine (B1), riboflavin (B2), nicotinamide (B3), pyridoxine (B6), folic acid (B9), and ascorbic acid (C), play important roles in cellular metabolism and redox-related processes. Following dietary intake, these vitamins are converted into their biologically active coenzyme forms, including thiamine pyrophosphate, flavin adenine dinucleotide, nicotinamide adenine dinucleotide, and pyridoxal-5′-phosphate, all of which are indispensable cofactors for enzymatic reactions supporting the production of ATP through central carbon and amino acid metabolisms [12], [13], [14]. In addition, ascorbic acid contributes to the maintenance of redox balance by scavenging reactive oxygen species and supporting redox-dependent metabolic reactions [15], [16].

In this context, a re-evaluation of legume crops — already widely cultivated as protein sources — from a micronutrient perspective, and particularly with respect to vitamin content, is warranted. The seeds and sprouts of mungbeans reportedly contain relatively high levels of vitamin C and folate compared with other major legume crops (USDA, 2010). Genotype-dependent variation in B1, B2, and B3 contents has also been observed in mungbean sprouts; however, these studies were limited by the inclusion of only three genotypes [17].

Mungbean sprouts consumed by humans are the product of germination, during which water uptake is followed by morphological changes, such as the emergence of the radicle and the development of the hypocotyl and shoot. Biochemical activities, including central metabolism, are reactivated concomitantly [18], [19]. This metabolic reactivation is often accompanied by changes in vitamin contents, and increased levels of vitamins during germination have been reported in various species of legumes [20], [21], [22].

Mungbean sprouts are widely consumed, and water-soluble vitamins are essential nutrients for human health; however, their contribution as dietary vitamin sources has not been sufficiently characterized. Previous studies have reported the presence of vitamins in mungbean sprouts, but information on vitamin composition and content among different mungbean genotypes remains limited. Antioxidant capacity is commonly used to evaluate the ability of food materials to suppress oxidative reactions, whereas tyrosinase inhibition serves as an enzyme-based indicator for assessing bioactivity-related properties [23], [24], [25], [26]. To obtain a comprehensive understanding of the nutritional and bioactivity-related characteristics of mungbean sprouts as consumed by humans, this study quantified six water-soluble vitamins (B1, B2, B3, B6, B9, and C) in sprouts derived from 34 genotypes of mungbeans and evaluated the antioxidant capacity and tyrosinase inhibition. The results provide foundational information that highlights the potential value of mungbean sprouts to serve as plant-based foods capable of improving efficient metabolic processes in the human body.

## 2 Materials and methods

### 2.1 Sample preparation

All mungbean seeds were harvested at the Gangneung-Wonju National University Experimental Farm in Gangneung, South Korea (37.77°N, 128.86°E) in 2021. The cultivation of mungbean sprouts followed a method suggested by [27], with some modifications. Fifty seeds were germinated for each of the 34 genotypes. The seeds were soaked in distilled water for 17 hours at 37 °C using an incubator (Jeiotech, ISS-4075R, Korea) for germination. The sprouted (germinated) seeds were cultivated for three days in a plant-growth chamber (Sundotcom, ST001A, Korea) at 28 ± 2 °C in the dark. During this cultivation period, water was sprayed automatically for 2 min every 4 h. The mungbean sprouts were harvested on the third day after germination, rapidly cooled with liquid nitrogen, and stored at −70 °C for 5 days. Subsequently, the sprouts were freeze-dried for 5 days using a freeze-dryer.

### 2.2 Extraction procedure

The freeze-dried samples were ground, weighed (0.1 g), and extracted in 70% ethanol under dark conditions for 24 h. The extract was then centrifuged at 15,000 rpm for 2 min, and the supernatant was filtered through a 0.22 µm syringe filter. Using this extract, 2,2′- azino-bis-3-ethylbenzthiazoline-6-sulphonic acid (ABTS) and 1,1-diphenyl-2-picrylhydrazyl (DPPH) assays were conducted to measure antioxidant capacity, while a tyrosinase inhibition assay was performed to assess the extract’s whitening activity.

#### 2.2.1 Vitamin B complex extracts in mungbean sprouts

The vitamin B complex standards used were thiamine hydrochloride (Chemfaces, CFN90068), riboflavin (Chemfaces, CFN90067), nicotinamide (Chemfaces, CFN99926), pyridoxine hydrochloride (Sigma, 58-56-0), and folic acid (Chemfaces, CFN98552). Thiamine, pyridoxine hydrochloride, and nicotinamide standards were prepared in a 0.1% solution of acetic acid containing 5 mM sodium hexanesulfonate, whereas riboflavin and folic acid standards were prepared in 0.1 N sodium hydroxide and subsequently diluted with 0.1% acetic acid containing 5 mM sodium hexanesulfonate. Standard calibration solutions were prepared over a concentration range of 0.5–8 μg/mL. For sample extraction, 0.1 g of the freeze-dried sample was mixed with 2 mL of a 0.1% acetic acid solution containing 5 mM sodium hexanesulfonate. After vortexing, the mixture was allowed to stand for 15 min and then sonicated for an additional 15 min. The extract was centrifuged at 15,000 rpm and −10 °C for 10 min, and the supernatant was filtered through a 0.22 μm syringe filter.

#### 2.2.2 Vitamin C extracts in mungbean sprouts

Ascorbic acid (Chemfaces, CFN90048) was used as the vitamin C standard. Standard and sample extractions were performed using metaphosphoric acid (Kanto Chemicals, 37267- 86-0). Ascorbic acid standard solutions were prepared at concentrations of 20, 40, 80, and 100 mg/L in 5% metaphosphoric acid. For sample extraction, 0.1 g of the sample was mixed with an equal volume of 10% metaphosphoric acid and allowed to stand for 10 min. Subsequently, 2 mL of 5% metaphosphoric acid was added and mixed thoroughly, followed by centrifugation at 15,000 rpm for 10 min. The supernatant was filtered through a 0.22 μm syringe filter

### 2.3 UPLC-photodiode array method

#### 2.3.1 Analysis of vitamin B complex by UPLC-PDA

Ultra performance liquid chromatography (UPLC) was performed for vitamin profiling (Nexera series equipped with MPM-40, SCL-40, SPD-M40, LC-40, SIL-40, and CTO-40 units from Shimadzu, Kyoto, Japan) using a photodiode array (PDA) detector. Vitamin B complexes and vitamin C were separated on a ZORBAX SB-C18 column (3.5 µm, 4.6 × 150 mm; Agilent, PN 863953-902, Santa Clara, USA). The mobile phase consisted of ultrapure water containing 0.1% acetic acid and 5 mM sodium hexanesulfonate (solvent A) and methanol containing 5 mM sodium hexanesulfonate (solvent B), delivered at a flow rate of 0.6 mL/min. The gradient program was as follows: 0–8 min, 80% A; 8–16 min, 80–40% A; 16–18 min, 40–20% A; 18–18.1 min, 20–0% A; 18.1–30 min, 0% A; 30.1 min, 0–80% A; and 30.1–35 min, 80% A. The oven temperature of the column was set at 30 °C, and the sample injection volume was 2 µL. The results were determined based on standard calibration curves (0.5–8 µg/mL) with three replicates. The photon wavelength of the detector scan range was set between 190 and 800 nm. The chromatograms of thiamine hydrochloride, riboflavin, nicotinamide, pyridoxine hydrochloride, and folic acid were extracted at 260, 270, 260, 290, and 280 nm respectively.

#### 2.3.2. Analysis of vitamin C by UPLC-PDA

The mobile phase consisted of 0.05 M potassium dihydrogen phosphate (solvent A) and acetonitrile (solvent B) mixed at a ratio of 60:40 (v/v) and was delivered at a flow rate of 0.2 mL/min. The column oven temperature was set at 40 °C, and the sample injection volume was 2 µL. Quantification was performed using standard calibration curves in the range of 20–100 mg/L, with three replicates. The chromatograms of ascorbic acid were extracted at 254 nm.

### 2.4 ABTS radical-scavenging assay

The ABTS radical-scavenging assay was measured according to a previously described protocol [28] with some modifications. The reaction between a 7 mM ABTS solution and 2.45 mM potassium persulphate in a 1:1 ratio resulted in the production of the ABTS radical cation. The ABTS solution was diluted with phosphate-buffered saline to reach an absorbance of 0.7 (±0.03) at 734 nm. Each sample (20 μL,1 mg/mL) was mixed with 180 μL of the ABTS solution in 96-well plates and the mixture was kept in the dark for 10 min. Absorbance at 738 nm was measured for each sample using a spectrophotometer. Ascorbic acid (FUJIFILM, 012–04802, Japan) (0, 1, 5, 10, 25, 50, and 100 mg/L) was used as the standard. The result was expressed as a percentage of scavenging achieved by the ABTS.

### 2.5 DPPH radical-scavenging assay

DPPH radical-scavenging activity was evaluated using an OxiTec DPPH Antioxidant Assay Kit (BIOMAX, BO-DPH-500, Korea) following the manufacturer’s instructions. Each sample solution (20 µL, 1 mg/mL) was mixed with assay buffer (80 µL) and DPPH radical solution (100 µL) in 96-well plates and incubated in the dark at room temperature for 30 min. The absorbance was measured at 517 nm using a spectrophotometer. Trolox (0, 40, 60, 80, and 100 mg/L) was used as a standard, and each standard (20 µL) was assayed under the same conditions as the samples. DPPH radical-scavenging activity was expressed as the percentage inhibition of the DPPH radical. All experiments were performed in triplicate.

### 2.6 Tyrosinase inhibitory activity assay

Tyrosinase inhibitory activity was determined using a modification of a previously reported dopachrome method [29] with 3,4-dihydroxy-L-phenylalanine (L-DOPA) as the substrate. A 70% ethanol mungbean extract was diluted to a final concentration of 5% ethanol for analysis. L-DOPA (Sigma-Aldrich, USA) was prepared in 0.1 M sodium phosphate buffer (pH 6.8; Biosesang, Republic of Korea). In each well of a 96-well microtiter plate, 25 μL of sample solution and 50 μL of mushroom tyrosinase solution (100 U) were mixed, followed by the addition of 25 μL of L-DOPA substrate, to initiate the reaction. The reaction mixture was incubated for 20 min at room temperature, and the absorbance was measured at 450 nm to determine dopachrome formation. A control containing 5% ethanol instead of the sample was used. Tyrosinase inhibitory activity was calculated as ([control − sample]/control) × 100.

### 2.7 Statistical analysis

Statistical analyses were performed in R software (R Core Team). Differences among groups were evaluated by one-way analysis of variance followed by a Duncan’s multiple range test. Pearson’s correlation analysis was also conducted. Statistical significance was set at p < 0.05.

## 3. Results

### 3.1 Vitamin composition and content in mungbean sprouts

The vitamin B complex (B1, B2, B3, B6, B9) and vitamin C contents were quantified in sprouts derived from 34 mungbean genotypes using a UPLC-PDA system (Figures 1 and 3). B1 content ranged from 2.68 ± 0.04 to 11.91 ± 0.16 mg/L, with genotype 21 (Betet) exhibiting the highest level and genotype 7 (JP99066) the lowest. B2 content ranged from 0.74 ± 0.01 to 1.76 ± 0.06 mg/L, with the maximum observed in genotype 5 (JP103138-1) and the minimum in genotype 1 (SM1409). B3 content varied from 5.85 ± 0.05 to 18.81 ± 0.18 mg/L, with genotype 20 (Arta ijo) showing the highest content and genotype 17 (Namwon, Jeollabuk-do, 1994-3237) the lowest. B6 content ranged from not detected to 2.76 ± 0.03 mg/L, with the highest level observed in genotype 24 (Dahyeon). B9 content ranged from 0.31 ± 0.01 to 1.44 ± 0.09 mg/L, with genotype 5 (JP103138-1) exhibiting the highest concentration and genotype 32 (Bohabe yellow mongo) the lowest. Vitamin C content ranged from 0.74 ± 0.14 to 15.58 ± 0.41 mg/L, with the lowest value detected in genotype 4 (JP229175) and the highest in genotype 19 (Acc. 363). All vitamins except B6 were detected across the analyzed genotypes, and B1 and B3 were the most abundant water-soluble vitamins. These results indicate genotype-dependent differences in vitamin composition and content.

**Figure 1.**
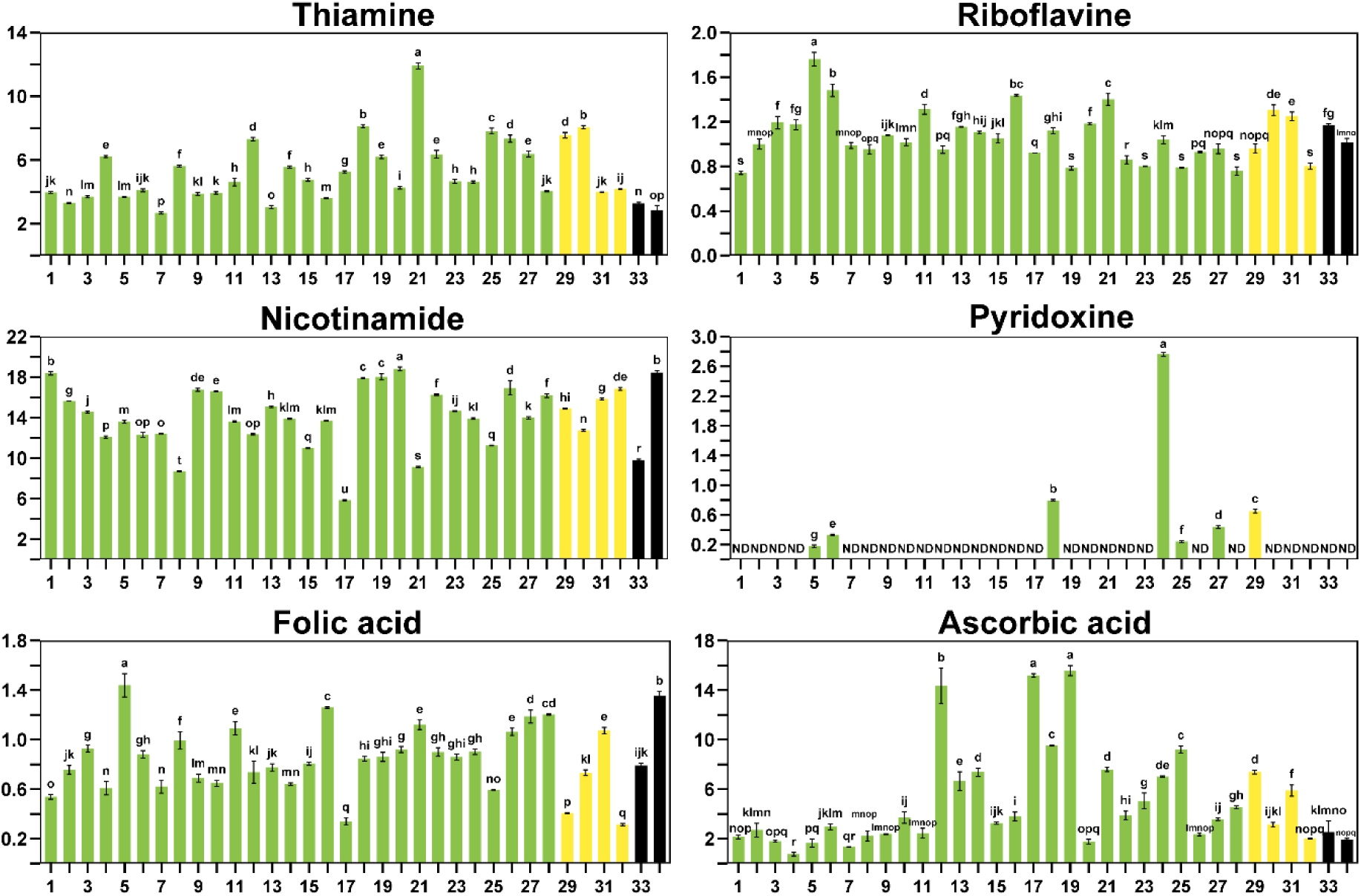
Contents of water-soluble vitamins, including the vitamin B complex (B1, B2, B3, B6, and B9) and vitamin C, in mungbean sprouts derived from 34 genotypes. The x and y axes indicate genotype and vitamin concentration (mg/L), respectively. Green, yellow, and black bars represent seed-coat color. Error bars indicate standard deviation. Different lowercase letters above the bars indicate statistically significant differences among genotypes (*P* < 0.05).

**Figure 2.**
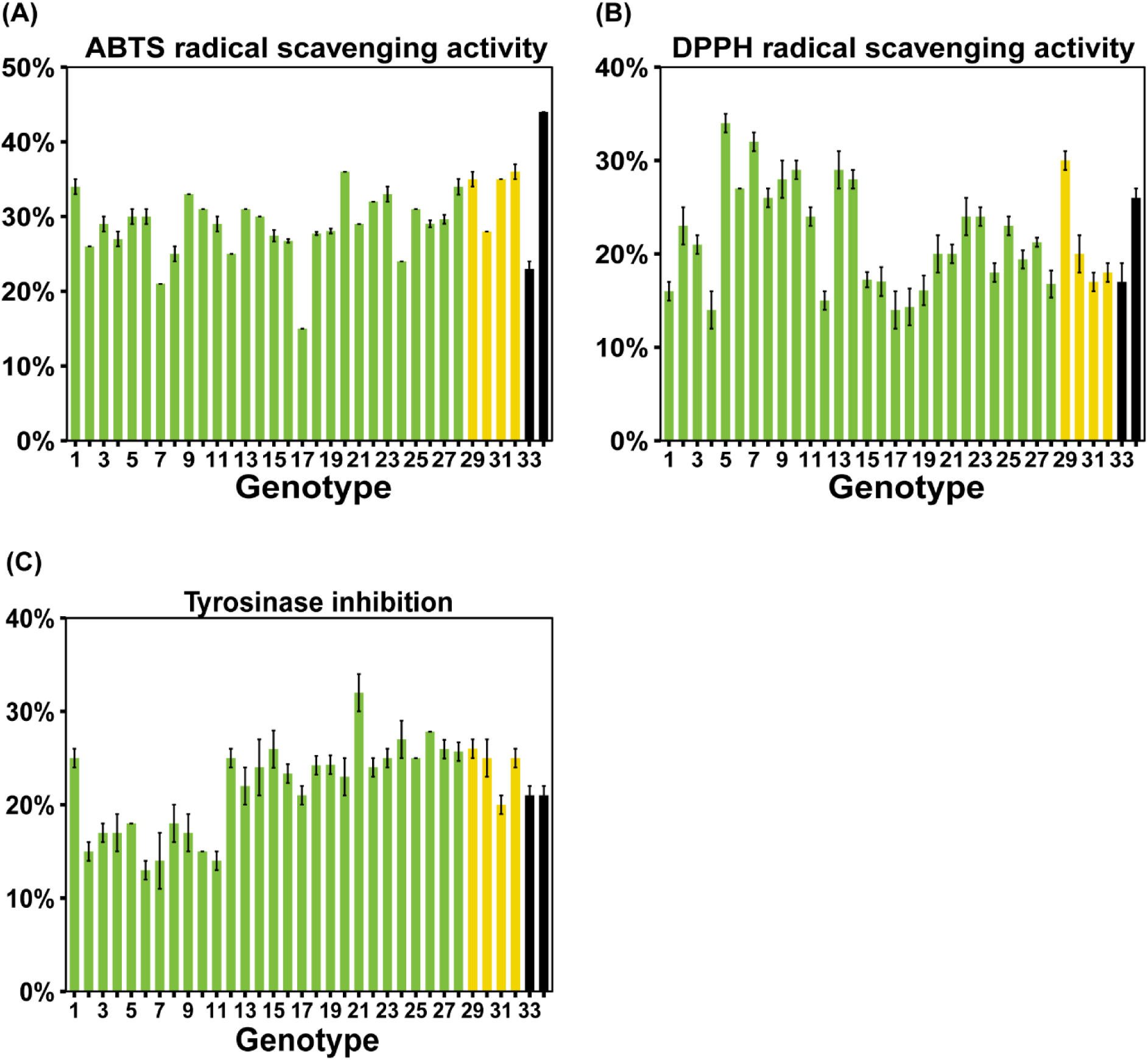
Antioxidant activity (ABTS and DPPH radical-scavenging activity) and tyrosinase inhibition of mungbean sprouts across 34 genotypes. The x and y axes indicate genotype and the percentage of ABTS and DPPH radical-scavenging activity or tyrosinase inhibition, respectively. Green, yellow, and black bars represent seed-coat color. Error bars indicate standard deviation. Different lowercase letters above the bars indicate statistically significant differences among genotypes (*P* < 0.05).

**Figure 3.**
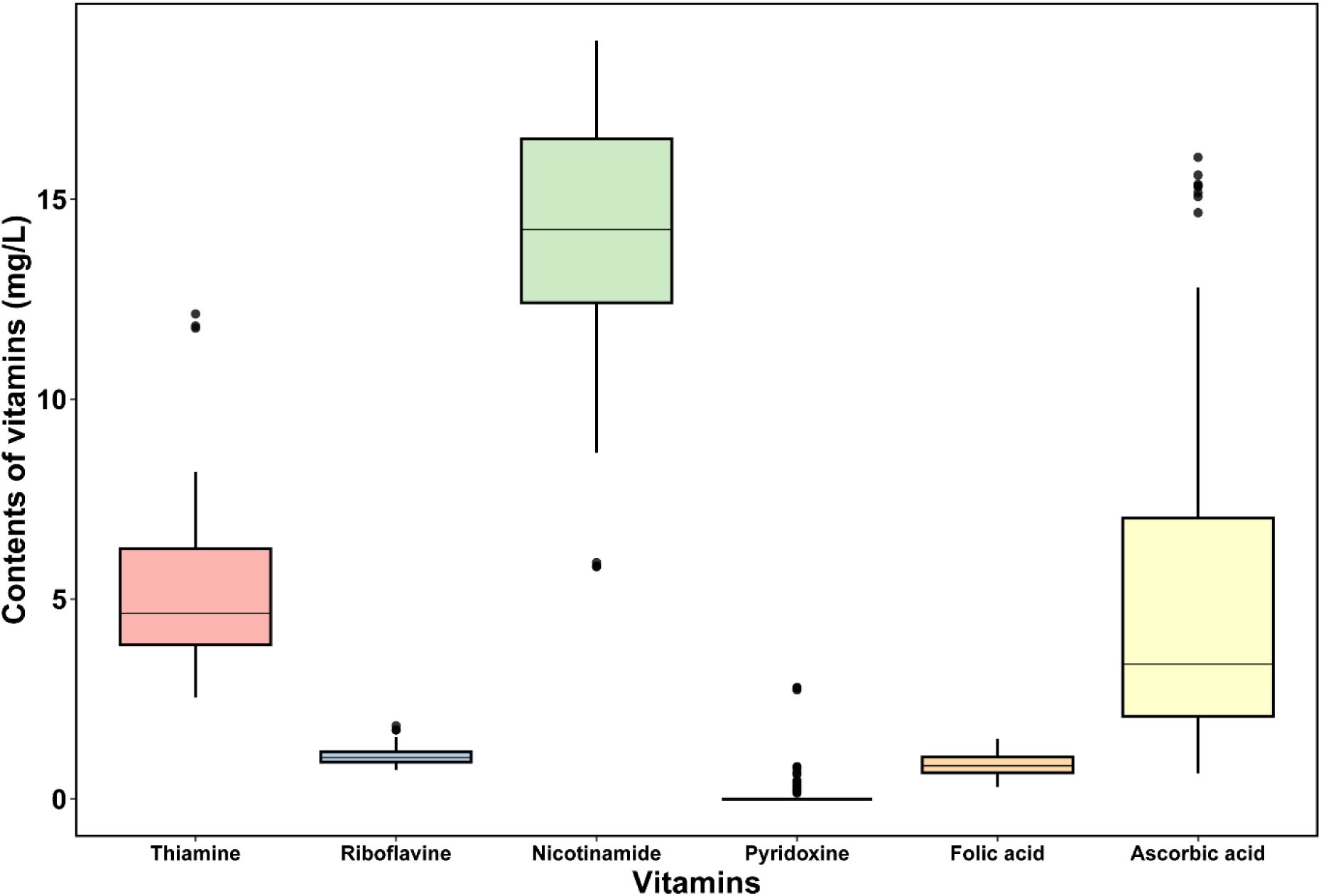
The vitamin B complex (B1, B2, B3, B6, B9) and vitamin C in mungbean sprouts of 34 genotypes. The x and y axes indicate vitamins and concentration of vitamins (mg/L), respectively.

### 3.2. Antioxidant capacity of mungbean sprouts

The antioxidant capacity of mungbean sprouts was evaluated across 34 genotypes using ABTS and DPPH radical-scavenging assays (Figure 2). ABTS radical-scavenging activity ranged from 15% ± 0% to 44% ± 0%, indicating substantial variation among genotypes. The lowest ABTS activity was observed in genotype 17 (Namwon, Jeollabuk-do, 1994-3237), and the highest activity was detected in genotype 34 (W190). DPPH radical-scavenging activity ranged from 14 ± 2% to 34 ± 1%. Genotype 5 (JP103138-1) exhibited the highest DPPH activity, while genotype 4 (JP229175) showed the lowest. Regardless of seed-coat color, antioxidant capacity varied markedly among genotypes.

### 3.3. Tyrosinase inhibition in mungbean sprouts

Tyrosinase is a key enzyme in melanin biosynthesis, catalyzing the hydroxylation of tyrosine to L-DOPA and the subsequent oxidation of L-DOPA to DOPA quinone, which ultimately leads to the formation of melanin. Inhibition of tyrosinase is therefore a widely applied strategy in the regulation of melanin production and skin pigmentation. Tyrosinase inhibition among mungbean sprout extracts ranged from 13% ± 1% to 32% ± 2%. The lowest inhibitory activity was observed in genotype 6 (JP103138-2), whereas genotype 21 (Betet) exhibited the highest. These results demonstrate clear genotype-dependent variation in tyrosinase inhibition, indicating that certain mungbean genotypes possess greater potential for suppressing melanin biosynthesis and may be valuable in food and cosmetic applications (Figure 2).

### 3.4 Correlation analysis among the contents of vitamins in mungbean genotypes

Pearson’s correlation analysis was conducted to examine relationships among the vitamin B complex (B1, B2, B3, B6, B9) and vitamin C content, antioxidant capacity (ABTS and DPPH assays), and tyrosinase inhibition across 34 mungbean genotypes (Figure 4). B3 content showed a positive correlation with ABTS radical-scavenging activity (r = 0.66). B2 content was positively correlated with B6 content (r = 0.46). B1 content showed a positive correlation with tyrosinase inhibition (r = 0.49). Overall, these findings suggest that variation in the composition and levels of specific vitamins may contribute to differences in antioxidant capacity and tyrosinase inhibitory activity.

**Figure 4.**
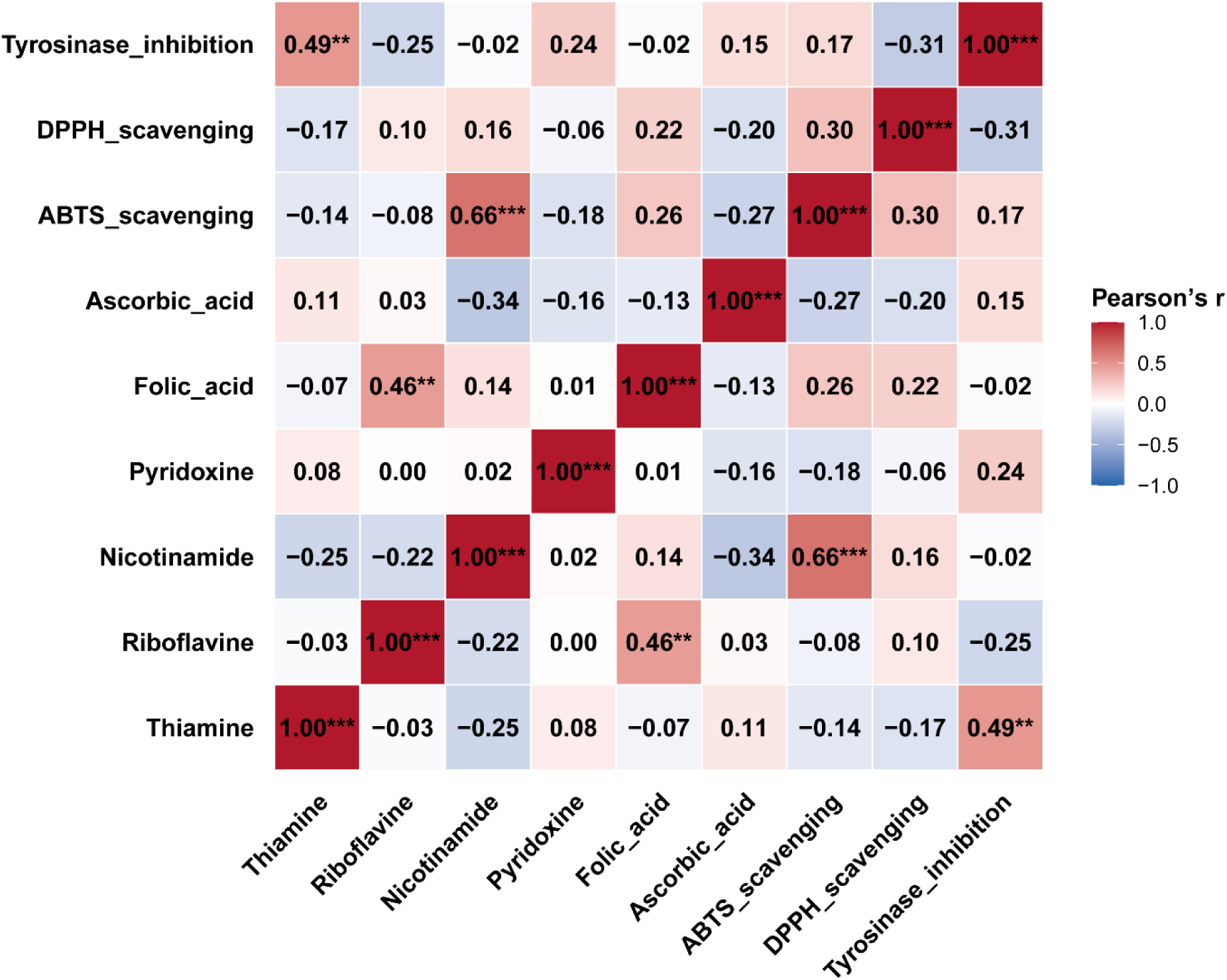
Correlation analysis among the contents of vitamins and antioxidants (ABTS and DPPH assays), as well as tyrosinase inhibition in mungbean sprouts. The correlations are presented using Pearson correlation coefficients and a color scale in which red indicates a positive correlation and blue indicates a negative correlation. *** *p* < 0.001, ** *p* < 0.01, * *p* < 0.05.

## Discussion

Micronutrient deficiencies remain a major global public health concern, particularly in developing countries. According to the World Health Organization, micronutrient deficiencies, including vitamin deficiencies, affect more than two billion people worldwide [30]. Because water-soluble vitamins cannot be synthesized endogenously in sufficient amounts and are readily excreted, continuous dietary intake is required to maintain adequate physiological levels. Previous studies have focused on one or a few individual vitamins using a limited number of genotypes. In contrast, the present study simultaneously evaluated six water-soluble vitamins (B1, B2, B3, B6, B9, and C) across 34 mungbean genotypes.

To assess the nutritional significance of vitamin contents in mungbean sprouts, vitamin contents were converted to a fresh-weight basis and compared with the 2024 U.S. Food and Drug Administration adult daily values (DV) [31]. A single 104 g serving of mungbean sprouts provides 46%–206% of the DV for B1 depending on genotype, indicating that sprouts from certain genotypes can meet or exceed the recommended daily intake of thiamine in a single serving. In contrast, B3 contributes 7%–24% of the DV, B2 up to 16%, B6 up to 33%, B9 up to 74%, and vitamin C up to 3% of the DV, representing relatively lower individual contributions compared with thiamine. However, energy metabolism proceeds efficiently when multiple vitamin-derived cofactors are present simultaneously. Because these cofactors participate in shared metabolic pathways, limiting a single vitamin can reduce overall metabolic efficiency even when other vitamins are present at sufficient levels [32], [33]. Taken together, mungbean sprouts provide a clear nutritional advantage through their high contribution of B1 while the concurrent presence of several B-complex vitamins suggests they can serve as a source capable of supporting efficient metabolic processes.

Beyond compositional assessment, the functional relevance of vitamin variation was evaluated using antioxidant activity assays. Cellular respiration inevitably generates reactive oxygen species, and several water-soluble vitamins contribute to redox regulation through their roles as cofactors in central metabolic pathways. In the present study, antioxidant activity assessed by ABTS assay differed significantly among mungbean sprout genotypes and showed a positive correlation with B3 content (r = 0.66***) (Figure 4). Given that B3 is a precursor of NAD⁺, a central cofactor in cellular redox metabolism, this association suggests that genotype-dependent variation in B3 accumulation may influence redox balance and antioxidant capacity following dietary intake. These findings indicate that the variation in vitamin composition and content among mungbean genotypes is reflected in differences in antioxidant activity, supporting their nutritional relevance.

In addition to redox-related antioxidant activity, B1 showed a significant correlation with tyrosinase-related activity (r = 0.49) (Figure 4), a parameter commonly used in the evaluation of whitening-related potential. In this regard, although vitamin C and B3 have been extensively studied [34], [35], whitening-related activity has also been suggested for certain other B vitamins, including B1. Previous in vitro studies using fungal models reported that B1 suppresses pigment formation by regulating intracellular signaling pathways, and patent-based reports have proposed that thiamine is an active component capable of exerting such effects [36]. Although the applicability of these mechanisms to human melanogenesis has not yet been fully established, the genotype-dependent correlations observed in the present study suggest that B1 is associated with functional activities beyond basic nutritional roles.

The present study revealed pronounced variation in vitamin content among mungbean genetic resources. Notably, concentrations of B1, the most abundant vitamin detected, ranged from 2.68 to 11.91 mg/L across genotypes, representing an approximately 4.5-fold difference, while B3 varied from 5.85 to 18.81 mg/L, showing a 3.2-fold range (Figures 1 and 3). These differences, observed under uniform growth conditions, indicate the presence of stable phenotypic variation in vitamin accumulation among mungbean accessions. Such variation is directly relevant to food quality, as it suggests that the nutritional value of mungbean sprouts consumed by end users can differ substantially depending on the genetic resources employed. In legume crops, including common beans, chickpeas, and soybeans, the importance of genetic resources as a foundation for nutritional improvement has been demonstrated through studies on seed vitamin traits [37], [38], [39]; however, this study is the first to evaluate the vitamin composition of mungbean sprouts across diverse genetic resources. The stable phenotypic variation identified in this study provides a basis for the genetic interpretation of vitamin-related traits and, ultimately, supports the selection of superior genetic resources for improving the nutritional quality of mungbean sprouts.

Although the present study revealed pronounced phenotypic variation in vitamin content among mungbean genetic resources, the genetic determinants underlying this variation were not directly examined. Evidence from model species nonetheless indicates that vitamin accumulation in plant-derived foods is under genetic control. In soybeans, mutation of the *GmPGL1* gene involved in B1 biosynthesis reduced levels of thiamine diphosphate, while disruption of the vitamin B6 biosynthesis gene *PDX1* in *Arabidopsis thaliana* (thale cress) resulted in decreased B6 production [40], [41]. These studies collectively indicate that specific biosynthetic genes play a key role in determining vitamin levels in edible plant tissues. Accordingly, the stable variation in vitamin accumulation observed in mungbean sprouts provides a basis for future genetic studies aimed at improving their nutritional quality. To date, mungbean research has focused primarily on agronomic traits such as yield and disease resistance, and few studies have addressed micronutrients derived from primary metabolism. In particular, comparative analyses of vitamin composition and content across diverse genetic resources are scarce. In the present study, substantial variation in vitamin content and composition was observed among mungbean genotypes. These findings provide a foundation for future research aimed at enhancing vitamin content and improving the nutritional quality of mungbeans.

## Conclusion

Mungbean sprouts are plant-based foods with meaningful qualities derived from their water-soluble vitamin content and associated functional properties. Genotype-dependent variation in vitamin content was observed across 34 genotypes under uniform germination conditions, indicating that the nutritional quality of mungbean sprouts can differ depending on the genetic resources used for food production. When vitamin contents were evaluated on a fresh-weight basis, thiamine and nicotinamide contributed substantially to daily dietary intake, supporting a role for mungbean sprouts in nutritionally relevant foods rather than as sources of trace micronutrients alone. In addition, differences in vitamin content were reflected in variation in antioxidant capacity and tyrosinase-related activity, indicating an association between vitamin levels and functional quality. Collectively, these findings provide a scientific basis for considering mungbean sprouts as value-added plant-based foods and for the selection and utilization of raw materials with superior nutritional profiles to ensure consistent nutritional quality in mungbean–based products.

## Data Availability

The data that support the findings of this study are available in the supplementary material of this article.

## Authors’ Contributions

S. Jeon performed the experiments and wrote the manuscript. H. Kwon revised the manuscript. S. Y. Song and S. D. Lim conducted statistical analysis and visualized the data. J. Ha designed the experiment, and revised the manuscript.

## Acknowledgments

This work was also supported by the National Research Foundation of Korea (NRF) grant funded by the Korea government (MSIT) (No. RS-2025-25431972).

This research was supported by the Regional Innovation System & Education (RISE) program through the Gangwon RISE Center, funded by the Ministry of Education (MOE) and the Gangwon State (G.S.), Republic of Korea. (2026-RISE-10-005)

This work was carried out with the support of “Research Program for Agriculture Science and Technology Development (Project No. PJ01738001)” Rural Development Administration, Republic of Korea.

## Supplementary Materials

This file contains Supplementary Tables S1-S2.

**Supplementary Table 1.** List of mungbean genotypes.

**Supplementary Table 2.** ABTS and DPPH radical-scavenging activities and tyrosinase inhibitory activity of mungbean genotypes.

