## Supplementary Tables for "Genotype-Dependent Variation in Vitamins B1, B2, B3, B6, B9, and C in Mungbean Sprouts"

**Supplementary Table 1.** List of mungbean genotypes.

| Number | Accession name | Origin | Number | Accession name | Origin |
|---|---|---|---|---|---|
| 1 | SM1409 | | 18 | V03720B-G | USA |
| 2 | JP229215 | CHN | 19 | Acc. 363 | PHL |
| 3 | JP229216 | CHN | 20 | Arta ijo | IDN |
| 4 | JP229175 | IND | 21 | Betet | IDN |
| 5 | JP103138-1 | PAK | 22 | PB-1 (benggolo Putih) | IDN |
| 6 | JP103138-2 | PAK | 23 | Fue Nutu | IDN |
| 7 | JP99066 | PAK | 24 | Dahyeon | KOR |
| 8 | JP229254 | IRN | 25 | Jeongdong, Sangju | KOR |
| 9 | CN900001 | THA | 26 | | |
| 10 | Suwon No.2 | KOR | 27 | Samgang | KOR |
| 11 | Gimje, Jeollabuk-do, 1985-3835 | KOR | 28 | VC1973A | TWN |
| 12 | Boeun, Chungcheongbuk-do, 1989-5499 | KOR | 29 | SM1501 | |
| 13 | Imsil, Jeollabuk-do, 1989-5600 | KOR | 30 | Yellowgram | |
| 14 | Vo3484 | PAK | 31 | ACC11 | PHL |
| 15 | Vo5551 | IRN | 32 | Bohabe yellow mongo | PHL |
| 16 | Collected Yecheon, 1992 | KOR | 33 | Damyang, Jeollanam-do, 1994-3231 | KOR |
| 17 | Namwon, Jeollabuk-do, 1994-3237 | KOR | 34 | W190 | |

*CHN, China; IND, India; PAK, Pakistan; IRN, Iran; THA, Thailand; KOR, Republic of Korea; USA, United States of America; PHL, Philippines; IDN, Indonesia; TWN, Taiwan.

**Supplementary Table 2.** ABTS and DPPH radical-scavenging activities and tyrosinase inhibitory activity of mungbean genotypes.

| Number | ABTS radical-scavenging activity | DPPH radical-scavenging activity | Tyrosinase inhibition |
|---|---|---|---|
| 1 | 34% ± 1%$^{cde}$ | 16% ± 1%$^{nopq}$ | 25% ± 1%$^{bcd}$ |
| 2 | 26% ± 0%$^{mn}$ | 23% ± 2%$^{gh}$ | 15% ± 1%$^{jklm}$ |
| 3 | 29% ± 1%$^{hij}$ | 21% ± 1%$^{hijk}$ | 17% ± 1%$^{ijkl}$ |
| 4 | 27% ± 1%$^{lm}$ | 14% ± 2%$^{q}$ | 17% ± 2%$^{ijk}$ |
| 5 | 30% ± 1%$^{ghi}$ | 34% ± 1%$^{a}$ | 18% ± 0%$^{hi}$ |
| 6 | 30% ± 1%$^{ghi}$ | 27% ± 0%$^{de}$ | 13% ± 1%$^{m}$ |
| 7 | 21% ± 0%$^{q}$ | 32% ± 1%$^{ab}$ | 14% ± 3%$^{lm}$ |
| 8 | 25% ± 1%$^{n}$ | 26% ± 1%$^{jklm}$ | 18% ± 2%$^{ij}$ |
| 9 | 33% ± 0%$^{e}$ | 28% ± 2%$^{cd}$ | 17% ± 2%$^{ijk}$ |
| 10 | 31% ± 0%$^{f}$ | 29% ± 1%$^{cd}$ | 15% ± 0%$^{klm}$ |
| 11 | 29% ± 1%$^{ghi}$ | 24% ± 1%$^{fgh}$ | 14% ± 1%$^{lm}$ |
| 12 | 25% ± 0%$^{no}$ | 15% ± 1%$^{opq}$ | 25% ± 1%$^{cde}$ |
| 13 | 31% ± 0%$^{fg}$ | 29% ± 2%$^{cd}$ | 22% ± 2%$^{efg}$ |
| 14 | 30% ± 0%$^{fgh}$ | 28% ± 1%$^{cd}$ | 24% ± 3%$^{de}$ |
| 15 | 27% ± 1%$^{kl}$ | 17% ± 1%$^{lmnop}$ | 26% ± 2%$^{bcd}$ |
| 16 | 27% ± 0%$^{lm}$ | 17% ± 2%$^{lmnopq}$ | 23% ± 1%$^{def}$ |
| 17 | 15% ± 0%$^{r}$ | 14% ± 2%$^{q}$ | 21% ± 1%$^{g}$ |
| 18 | 28% ± 0%$^{jkl}$ | 14% ± 2%$^{pq}$ | 24% ± 1%$^{cde}$ |
| 19 | 28% ± 0%$^{ijk}$ | 16% ± 2%$^{nopq}$ | 24% ± 1%$^{cde}$ |
| 20 | 36% ± 0%$^{kl}$ | 20% ± 2%$^{ijkl}$ | 23% ± 2%$^{bcd}$ |
| 21 | 29% ± 0%$^{hij}$ | 20% ± 1%$^{jklm}$ | 32% ± 2%$^{a}$ |
| 22 | 32% ± 0%$^{fg}$ | 24% ± 2%$^{fgh}$ | 24% ± 1%$^{cde}$ |
| 23 | 33% ± 1%$^{de}$ | 24% ± 1%$^{efg}$ | 25% ± 1%$^{bcd}$ |
| 24 | 24% ± 0%$^{op}$ | 18% ± 1%$^{klmn}$ | 27% ± 2%$^{bc}$ |
| 25 | 31% ± 0%$^{f}$ | 23% ± 1%$^{ghi}$ | 25% ± 0%$^{bcd}$ |
| 26 | 29% ± 0%$^{ij}$ | 19% ± 1%$^{jklm}$ | 28% ± 0%$^{b}$ |
| 27 | 30% ± 1%$^{ghi}$ | 21% ± 0%$^{hij}$ | 26% ± 1%$^{bcd}$ |
| 28 | 34% ± 1%$^{cde}$ | 17% ± 1%$^{mnopq}$ | 26% ± 1%$^{bcd}$ |
| 29 | 35% ± 1%$^{bc}$ | 30% ± 1%$^{bc}$ | 26% ± 1%$^{bcd}$ |
| 30 | 28% ± 0%$^{ijk}$ | 20% ± 2%$^{hijk}$ | 25% ± 2%$^{bcd}$ |
| 31 | 35% ± 0%$^{bcd}$ | 17% ± 1%$^{mnopq}$ | 20% ± 1%$^{gh}$ |

| | | | |
|---|---|---|---|
| 32 | 36% ± 1%[b] | 18% ± 1%[klmno] | 25% ± 1%[cd] |
| 33 | 23% ± 1%[p] | 17% ± 2%[lmnopq] | 21% ± 1%[fg] |
| 34 | 44% ± 0%[a] | 26% ± 1%[def] | 21% ± 1%[fg] |
